# A maternal-effect *Padi6* variant results in abnormal nuclear localization of DNMT1 and failure of epigenetic reprogramming and zygotic genome activation in mouse embryos

**DOI:** 10.1101/2023.10.09.561545

**Authors:** Carlo Giaccari, Francesco Cecere, Lucia Argenziano, Antonio Galvao, Dario Acampora, Gianna Rossi, Bruno Hay Mele, Maria Vittoria Cubellis, Flavia Cerrato, Simon Andrews, Sandra Cecconi, Gavin Kelsey, Andrea Riccio

## Abstract

PADI6 belongs to the multi-protein sub-cortical maternal complex (SCMC) that is present specifically in mammalian oocytes and early embryos. Maternal inactivation of SCMC genes generally results in early embryo lethality. In humans, variants in a subset of SCMC genes have been found in the healthy mothers of children affected by genomic imprinting disorders and characterized by multi-locus imprinting disturbances (MLID). However, how the SCMC controls the DNA methylation required to regulate imprinting remains poorly defined. To address this issue, we generated a mouse line carrying a *Padi6* missense variant that had been identified in the mother of two sisters affected by Beckwith-Wiedemann syndrome and MLID. We found that if homozygous in female mice this variant resulted in interruption of embryo development at the 2-cell stage. Single-cell DNA methylation and RNA analyses demonstrated genomic hypermethylation, down-regulation of zygotic genome activation (ZGA) genes and up-regulation of maternal decay genes in 2-cell embryos from homozygous females. In addition, immunofluorescence analysis showed abnormal localization of DNMT1 and UHRF1 in mutant oocytes and zygotes. Taken together, this study demonstrates that PADI6 controls the subcellular localization of DNMT1 that is necessary for pre-implantation epigenetic reprogramming and ZGA.

## Introduction

DNA methylation inherited from the gametes is substantially reprogrammed during the oocyte-to-embryo transition (Sendžikaitė and Kelsey 2019). After fertilization, embryonic cells lose oocyte-and sperm-specific DNA methylation and regain totipotency. During pre-implantation development, paternally-inherited DNA is rapidly demethylated, at least in part involving activity of Ten-eleven translocation (TET) enzymes, while maternally-inherited DNA methylation is lost in a passive manner during cellular proliferation through nuclear exclusion of DNMT1 and its accessory protein UHRF1 (Chen and Zhang 2020). Set against this global trend, imprinted genes need to retain their gamete-of-origin DNA methylation (Barlow and Bartolomei 2014). Immediately after fertilization, the zygotic genome is transcriptionally silent and early embryo development is controlled by maternal mRNAs and proteins (Aoki 2022). Zygotic genome activation (ZGA) in mice is initiated between the mid-1-cell and early 2-cell stages (minor ZGA) and proceeds during the mid-to-late 2-cell stage (major ZGA) (Aoki 2022).

In mammals, the SCMC is a multi-protein complex that has been associated with multiple biological functions occurring during the oocyte to embryo transition (Li et al. 2008; Bebbere et al. 2021). A number of proteins have been demonstrated or proposed to be part of the SCMC, including KHDC3L, OOEP, PADI6, TLE6, ZBED3, as well as several NLRP family members (NLRP2, NLRP4f, NLRP5, NLRP9a/b/c and, in humans NLRP7) (Bebbere et al. 2021). All SCMC proteins are encoded by maternal-effect genes. PADI6 is an oocyte-specific protein belonging to the peptidylarginine deiminase (PAD) protein family but without evident enzymatic activity in vitro (Raijmakers et al. 2007; Williams and Walport 2023). Maternal PADI6 is required for embryo development beyond the 2-cell stage and formation of the oocyte lattices that are cytoplasmic structures believed to work as ribosomal storage for the early embryo (Esposito et al. 2007). Maternal *Padi6*-mutant 2-cell embryos have ribosomal components and *de novo* protein synthesis impaired, as well as drastically reduced transcription levels (Yurttas et al. 2008). In humans, biallelic loss-of-function mutations of *PADI6* are associated with female infertility and hydatidiform mole (Qian et al. 2018). Furthermore, both biallelic and monoallelic *PADI6* variants have been found in the healthy mothers of children affected by genomic imprinting disorders (Eggermann et al. 2022; Williams and Walport 2023).

Genomic imprinting is the gamete of origin-dependent epigenetic marking and expression of genes, and is required for normal development (Barlow and Bartolomei 2014). In humans, its deregulation affects growth, metabolism and neurological functions (Eggermann et al. 2023). Imprinting disorders are associated with single or multi-locus DNA methylation abnormalities, which in turn can be traced back to genetic variants occurring in *cis* or in *trans* (Monk et al. 2019). In particular, maternal-effect SCMC gene variants are associated with multi-locus imprinting disturbance (MLID, see (Eggermann et al. 2022). Very little is known about the mechanisms linking the SCMC and DNA methylation. So far, it has been demonstrated that a hypomorphic *KHDC3L* variant results in globally impaired *de novo* methylation in human oocytes (Demond et al. 2019), while *Nlrp2* knockout in mice leads to abnormal subcellular localization of DNMT1 in oocytes and DNA methylation abnormalities in mouse embryos (Mahadevan et al. 2017; Yan et al. 2023) and impacts expression of the histone Demethylase KDM1B (Anvar et al. 2023). More recently, it has been shown that the maternal knockout of a further but not SCMC-associated *Nlrp*-gene (*Nlrp14*) also results in altered localization of DNMT1 and genomic hypermethylation in early embryos (Yan et al. 2023).

Since inactivation of SCMC genes generally leads to early embryo lethality, studying the mechanisms by which these maternal-effect genes control maternal to zygotic transition in mammals has been challenging. Here, we describe a mouse line carrying a variant of the *PADI6* gene found in the genome of the mother of two sibs affected by Beckwith-Wiedemann syndrome (BWS) and MLID (Cubellis et al. 2020). By applying a method of combined single-cell profiling of gene expression and DNA methylation, we investigated the molecular changes occurring in early development. We found that, if present in the mother, this variant severely affects early mouse development. Epigenetic reprogramming and ZGA were prevented in the mutant 2-cell embryos concomitant with the nuclear mis-localization of DNMT1 and UHRF1.

## Results

### Generation of the *Padi6*^P620A^ mouse line

To study the role of maternal-effect *Padi6* variants on DNA methylation programming and imprinting, we generated a knock-in mouse line modelling the human missense mutation P632A that was found in compound heterozygosity with a truncating mutation in a family with two siblings affected by Beckwith-Wiedemann syndrome and MLID (Cubellis et al. 2020). Proline 632 is located in a highly conserved region of the PADI6 protein and corresponds to residue 620 of the mouse protein (Fig. 1a). So, the variant chr4, g.140727767 C → G; P620A of *Padi6* exon 16 was introduced into the genome of mouse embryonic stem (ES) cells by homologous recombination through the use of a FLP/frt system vector (Fig. 1b). The presence of the mutation in ES cell clones was demonstrated by Sanger sequencing and Southern blotting analysis (Fig. S1a,b). Chimeric mice were then obtained by injection of the positive clones into mouse blastocysts and germline transmission was confirmed by PCR. To remove the NEO cassette, the knock-in mouse line was crossed with a line carrying the *Flp* recombinase gene, and excision was confirmed by PCR (Fig. S1c). To exclude any possible rearrangement during recombination, we determined the whole-genome sequence of somatic DNA of a *Padi6*^P620A/P620A^ female mouse. We confirmed the presence of the chr4, g.140727767 C → G variant and demonstrated the absence of any CNV or structural variant involving the *Padi6* gene (Fig. S1d and data not shown).

**Figure 1.**
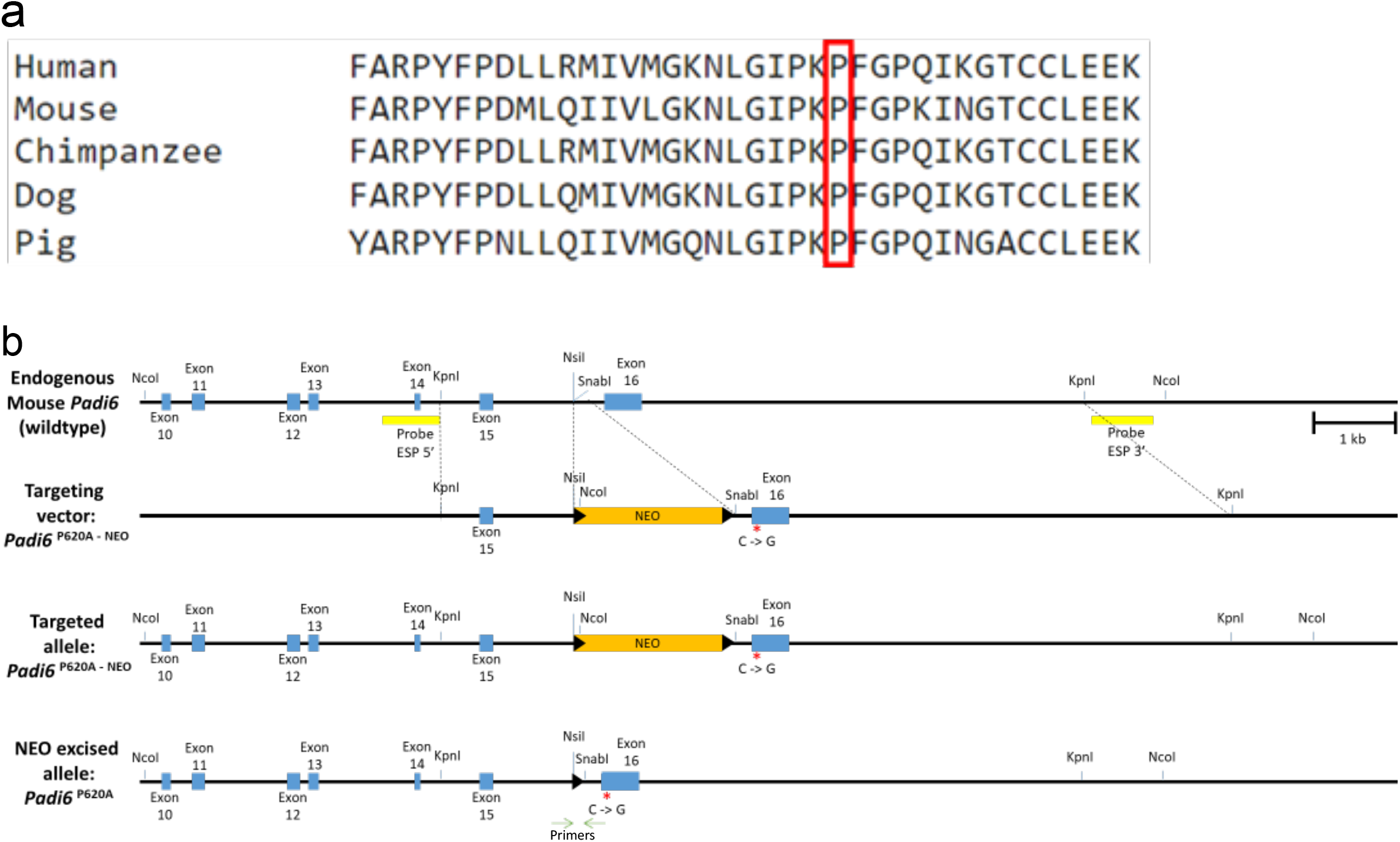
Generation of the *Padi6*^P620A^ allele. (**a**) Multiple Sequence Alignment of PADI6 Orthologous Proteins from different species. The red rectangle identifies the site of interest. (**b**) From the top: the mouse *Padi6* endogenous locus, the targeting vector (*Padi6*^P620A-NEO^), the correctly targeted allele with the neomycin resistance cassette (*Padi6*^P620A-NEO^) and the targeted allele after NEO excision (*Padi6*^P620A^). The blue rectangles depict the *Padi6* exons. The orange rectangle and the black triangle depict the NEO cassette and the *frt* sites, respectively. The mutation is marked by a red asterisk. The probes (ESP 5’ and ESP3’) used for Southern blot screening are indicated by yellow lines below the endogenous locus and the PCR primers used for confirming targeted allele and NEO excision are indicated by green arrows.

### Phenotypic characterization of the *Padi6*^P620A^ mouse line

From heterozygous *Padi6*^P620A/+^ inter-breeding, we obtained 12 litters including 23 *Padi6*^+/+^, 40 *Padi6*^P620A/+^ and 26 *Padi6*^P620A/P620A^ pups, a ratio that did not significantly differ from the expected genotype distribution (Χ^2^ = 0.6192, *p* = 0.4313) (Fig. 2a). Both *Padi6*^P620A/+^ and *Padi6*^P620A/P620A^ mice were viable, did not display any visible morphological anomaly, grew normally and with normal lifespan. Similar to *Padi6*^+/+^ animals, all follicular stages and corpora lutea were observed in the ovaries of 4-8 week-old *Padi6*^P620A/+^ and *Padi6*^P620A/P620A^ females (Fig. 2b,c and data not shown). Similar level of *Padi6* mRNA was demonstrated in MII oocytes of the *Padi6*^+/+^ and *Padi6*^P620A/P620A^ mice by single-cell RNA sequencing (scRNA-seq) analysis (Fig. 2d). However, while the PADI6 protein was strongly detected in the ovary and correctly localized in the oocyte cytoplasm of the control mice (Kim et al. 2010), we found that its level was decreased in the *Padi6*^P620A/P620A^ mice below the detection levels of western blotting and immunofluorescence (Fig. 2e-g). Then, we tested whether the *Padi6* P620A variant affected mouse fertility. We obtained litters of similar size from natural matings between 2-4 month-old *Padi6*^+/+^ and *Padi6*^P620A/+^ females with *Padi6*^+/+^ males. In contrast, *Padi6*^P620A/P620A^ females were completely unable to have pups (Fig. 2h). We performed a similar study on male fertility, but no significant difference was observed in litter size from 2-4 month-old *Padi6*^+/+^, *Padi6*^P620A/+^ or *Padi6*^P620A/P620A^ males crossed with *Padi6*^+/+^ females (Fig. 2i). Thus, the P620A variant causes a strong reduction of the PADI6 protein level and profoundly affects female fertility but had no effect on male fertility.

**Figure 2.**
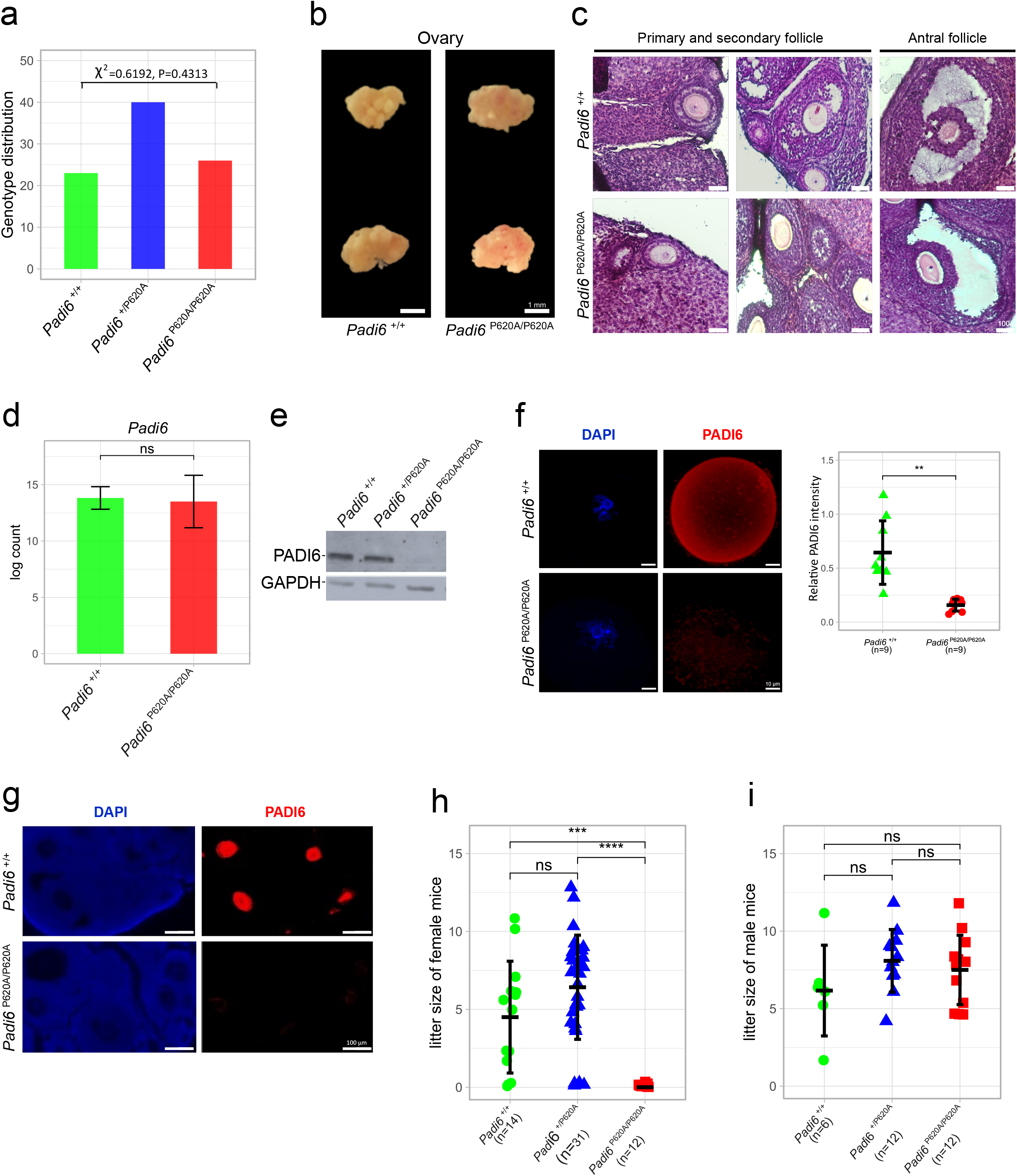
Characterization of the mouse line carrying the *Padi6*^P620A^ mutation. (**a**) Genotype distribution of the offspring derived from *Padi6*^P620A/+^ matings. (**b**) Comparison of morphological characteristics between *Padi6*^+/+^ and *Padi6*^P620A/P620A^ ovaries. The mice were derived from the same litter and the ovaries were collected from 8 week-old females. (**c**) Hematoxylin and eosin staining on paraffin-embedded *Padi6*^+/+^ (top) and *Padi6*^P620A/P620A^ (bottom) ovaries; the left and central panels show primary and secondary follicles, the right panel shows an antral follicle. (**d**) Barplot representing the *Padi6* expression level calculated from log2 mean count of scRNAseq data of 18 *Padi6*^+/+^ (green) and 20 *Padi6*^P620A/P620A^ (red) MII oocytes. (**e**) Western blotting analysis of PADI6 and GAPDH in *Padi6*^+/+^, *Padi6*^+/P620A^ and *Padi6*^P620A/P620A^ ovaries. The data shown are representatives of three independent experiments. (**f**) Left, representative images of immunostaining of *Padi6*^+/+^ (top) and *Padi6*^P620A/P620A^ (bottom) GV oocytes with anti-PADI6 antibodies (red) and DAPI (blue). Right, quantification of PADI6 immunostaining intensity. *n* denotes the number of oocytes analysed in two independent experiments. (**g**) Representative images of paraffin-embedded ovarian sections from 8 week-old *Padi6*^+/+^ (top) and *Padi6*^P620A/P620A^ (bottom) mice immunostained with anti-PADI6 antibodies (red) and DAPI (blue). Consistent results were obtained in two independent experiments. (**h**) Female and (**i**) male fertility of *Padi6*^+/+^ (green), *Padi6*^P620A/+^ (blue) and *Padi6*^P620A/P620A^ (red) mice; 2-4 month old females (**h**) and males (**i**) of the indicated genotypes were mated with *Padi6*^+/+^ males and females, respectively. *n* denotes the number of mice analysed. The data shown in **d**, **f**, **h** and **i** are mean ± s.d. and were analysed using an unpaired two-tailed Student’s *t*-test (ns, *P* < 0.05; \*\**P* < 0.01; \*\*\**P* < 0.001).

### The *Padi6*^P620A/P620A^ oocytes show altered subcellular localization but normal expression of SCMC components

To study the effect of the *Padi6*^P620A^ variant on the SCMC we determined the expression of the other components of the complex in oocytes. qRT-PCR and scRNA-seq analyses revealed no significant difference in the expression of the *Ooep*, *Nlrp5* and *Tle6* genes among *Padi6*^+/+^, *Padi6*^P620A/+^ and *Padi6*^P620A/P620A^ oocytes (Fig. 3a,b). Similarly, comparable NLRP5 and TLE6 protein levels were detected in *Padi6*^+/+^, *Padi6*^P620A/+^ and *Padi6*^P620A/P620A^ ovaries by western blot analysis (Fig. 3c). However, different results were obtained by immunofluorescence (Fig. 3d). While typical subcortical localization of OOEP was observed in the germinal vesicle (GV) oocytes of *Padi6*^+/+^ mice, reduced cytoplasmic fluorescence and increased signal in the nucleus were found in the *Padi6*^P620A/P620A^ mutants (Fig. 3d). These results indicate that the *Padi6*^P620A^ variant does not affect the expression of the SCMC components other than PADI6 but modifies their subcellular localization.

**Figure 3.**
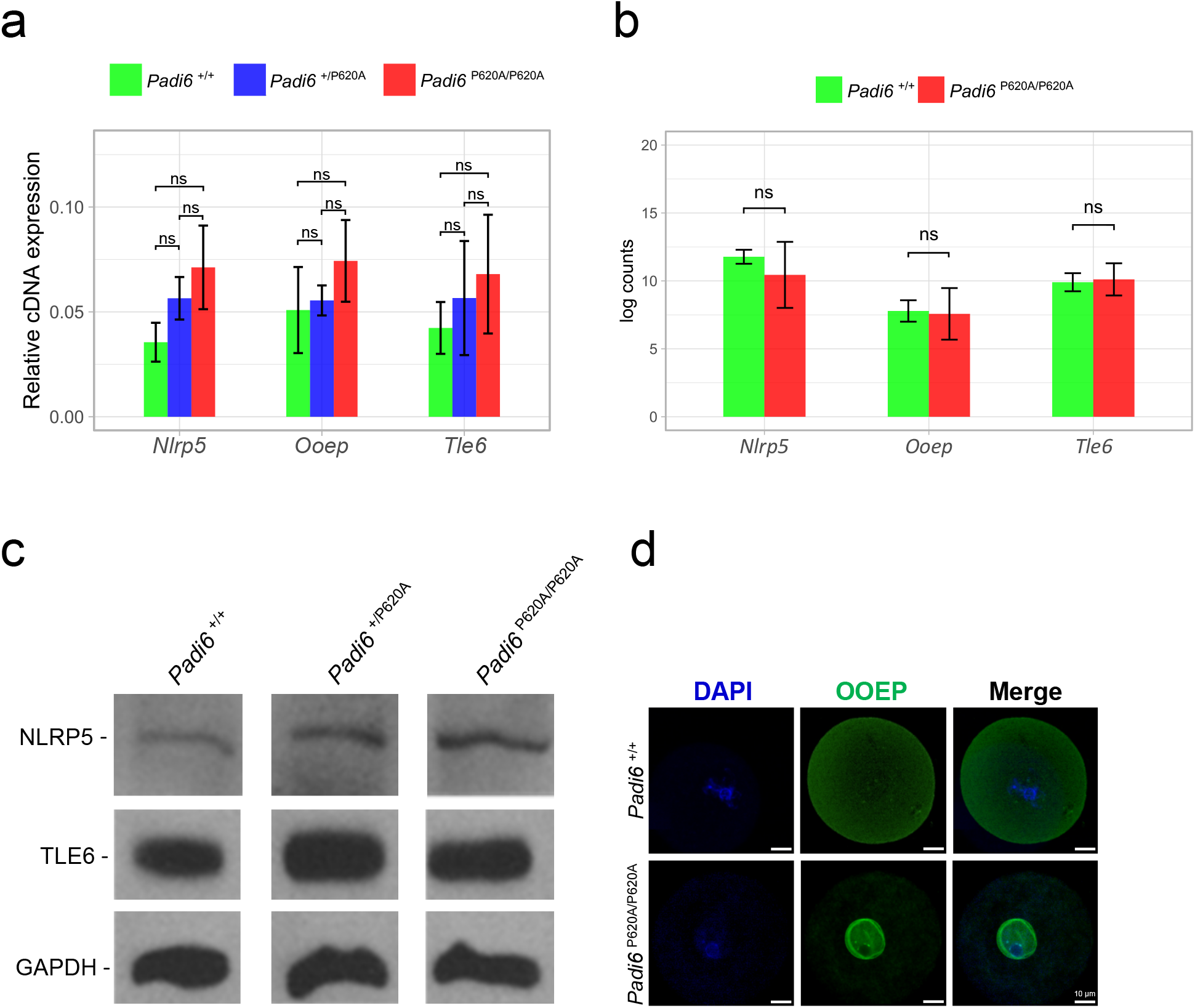
Effect of the *Padi6*^P620A^ variant on gene expression and protein level of the components of the SCMC. (**a**) *Nlrp5*, *Ooep* and *Tle6* gene expression analysis in *Padi6*^+/+^ (green), *Padi6*^P620A/+^(blue) and *Padi6*^P620A/P620A^ (red) ovaries, as assessed by RT-qPCR. The histograms show the average gene expression levels obtained in three independent experiments, after normalization against the level of *β-actin*. (**b**) Histograms representing the *Nlrp5*, *Ooep* and *Tle6* expression levels calculated with log2 mean counts of scRNAseq data of 18 *Padi6*^+/+^ (green) and 20 *Padi6*^P620A/P620A^ (red) MII oocytes. (**c**) Western blot analysis of NLRP5, TLE6 and GAPDH in *Padi6*^+/+^, *Padi6*^+/P620A^ and *Padi6*^P620A/P620A^ ovaries. The data shown are representatives of three independent experiments. (**d**) Representative images of immunostaining of *Padi6*^+/+^ (top) and *Padi6*^P620A/P620A^ (bottom) GV oocytes with anti-OOEP antibodies (green) and DAPI (blue). Consistent results were obtained in three independent experiments. The data shown in **a** and **b** are mean ± s.d. and were analysed using an unpaired two-tailed Student’s *t*-test (ns, *P* < 0.05; \*\**P* < 0.01; \*\*\**P*. < 0.001).

### Developmental failure and abnormal cell divisions of *Padi6*^MatP620A/+^ embryos

To investigate if the infertility of *Padi6*^P620A/P620A^ female mice was caused by embryonic death,we performed in vitro fertilization (IVF) experiments with ovulated MII oocytes collected from *Padi6*^+/+^ and *Padi6*^P620A/P620A^ females. After hormonal treatment, no difference in in the number of ovulated eggs was observed between *Padi6*^P620A/P620A^ and *Padi6*^+/+^ females (Fig. 4a). Also, both genotypes of mice showed a success rate of fertilisation higher than 50% (65% and 55% of 2-cell embryos for *Padi6*^+/+^ and *Padi6*^P620A/P620A^ oocytes, respectively) 24 hours after the start of IVF (Fig. 4b). Greater differences between the two groups began to appear 48 hours following IVF (Fig. 4c,d). While all the *Padi6*^+/+^ 2-cell embryos continued to develop normally, only 24% of the embryos derived from *Padi6*^P620A/P620A^ females (hereafter referred to as *Padi6*^MatP620A/+^ embryos) were able to reach the 4-cell stage. After 72 hours, most of the control embryos had reached the morula stage (95%). In contrast, the *Padi6*^MatP620A/+^ embryos showed significant impairment of embryonic development, with the majority blocked at 2-or 4-cell stages and a small percentage at 8-cell stage, and many displaying reduced number of blastomers and/or morphologic abnormalities and/or asymmetric cell divisions (Fig. 4c,d). Thus, after 3 days of development, almost all the *Padi6*^MatP620A/+^ embryos were either degenerated or arrested. After 96 hours, control embryos reached the blastocyst stage, while only a small percentage of *Padi6*^MatP620A/+^ embryos showed a degree of compaction similar to morula (Fig. 4d). More than 120 hours from IVF, very few *Padi6*^MatP620A/+^ embryos with a blastocyst-like morphology were found (Fig. 4e). In summary, the *Padi6*^P620A^ variant displayed a maternal-effect phenotype with abnormal embryonic development and mostly interrupted at the 2-cell stage, if present in homozygosity.

**Figure 4.**
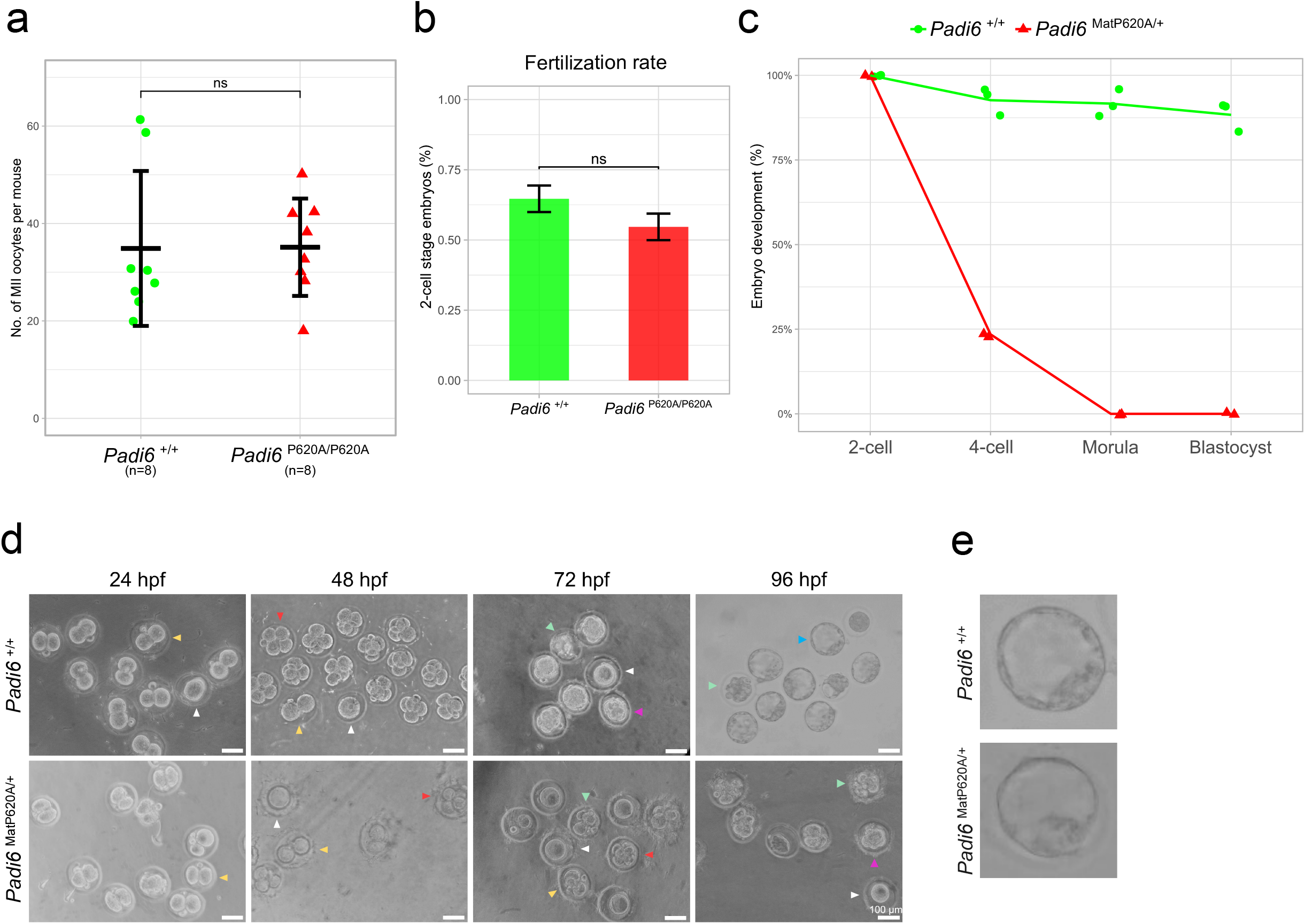
Effect of *Padi6*^P620A^ variant on IVF and embryo development. (**a**) Number of MII oocytes ovulated per female mouse after PMSG + hCG hormone treatment. *n* denotes the number of females analysed. (**b**) Bar plot representing the fertilization rate after IVF of MII oocytes from 8 week-old *Padi6*^+/+^ (green) and *Padi6*^P620A/P620A^ (red) females with wild-type sperm. (**c**) Line plot showing the percentage of *Padi6*^+/+^ (green) and *Padi6*^MatP620A/+^ (red) embryo developed after IVF. Each dot and triangle represent independent experiments. (**d**) Representative still frames of *Padi6*^+/+^ (top) and *Padi6*^MatP620A/+^ (bottom) embryos at 24, 48, 72 and 96 hours after IVF. Arrows indicate different developmental stages: unfertilized oocyte (white), 2-cell embryo (yellow), 4-cell embryo (red), morula (purple), blastocyst (light blue) and abnormal embryo (light green). (**e**) Representative example of *Padi6*^+/+^ (top) and *Padi6*^MatP620A/+^ (bottom) blastocyst. The data shown in **a** and **b** are mean ± s.d. and were analysed using an unpaired two-tailed Student’s *t*-test (ns, *P* < 0.05; \*\**P* < 0.01; \*\*\**P* < 0.001).

### The *Padi6*^P620A^ variant leads to transcriptome differences indicative of defective oocyte maturation

To look for the effect of the *Padi6*^P620A^ variant on the oocyte transcriptome, we analysed MII oocytes by scRNA-seq. Comparison of the scRNA-seq datasets of the *Padi6*^+/+^ and *Padi6*^P620A/P620A^ oocytes by global principal component analysis (PCA) resulted in some degree of separation along Principal Component 1 (PC1), particularly for the *Padi6*^P620A/P620A^ cells (Fig. 5a). 706 genes were detected as being differentially expressed (|log2FC| >= 1 and FDR < 0.01) and defined as MII-DEGs (Supp. Table S1). Out of these, 117 were down-regulated and 589 up-regulated (Fig. 5b). Using the MII-DEGs, three different clusters of oocytes were identified by unsupervised hierarchical clustering of scRNA-seq data (Fig. 5c). Clusters 1 and 2 corresponded to most of the *Padi6*^P620A/P620A^ oocytes and cluster 3 to most *Padi6*^+/+^ oocytes, thus demonstrating two groups of *Padi6*^P620A/P620A^ oocytes according to their RNA profiles. In particular, a subset of DEGs (n=345) was highly abundant in Cluster 2 but showed heterogeneous expression in Cluster 1 (Fig. 5a, Supp. Table S2). By looking at the function of these genes, we found that 17% of them (n=60) were reported as GV oocyte markers (Zhao, 2020) (Fig. 5c,d, Supp. Table S3) and associated with the Gene Ontology terms GO:BP system development (40%) and GO:CC cytoplasm (75%) (Fig. 5e). Although the cells were morphologically indistinguishable, these results suggest the existence of different stages of maturation within the pool of *Padi6*^P620A/P620A^ MII oocytes analysed by scRNA-seq.

**Figure 5.**
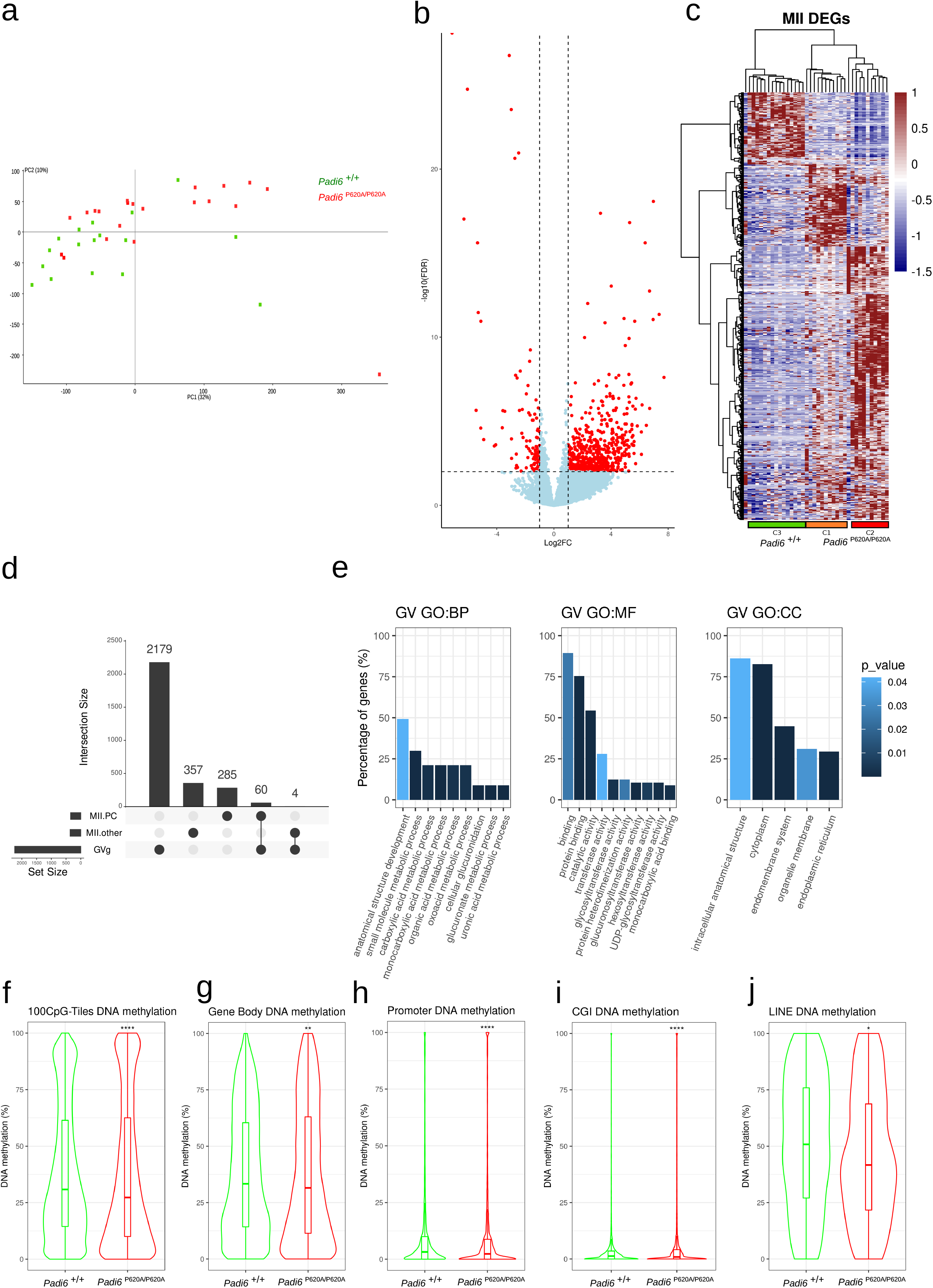
scRNA-seq and scBS-seq analyses of *Padi6*^P620A/P620A^ and *Padi6*^+/+^ MII oocytes. (**a**) PCA of all the genes covered by the scRNAseq experiment. The Principal Component 1 (PC1) is plotted on the x-axis and the Principal Component 2 (PC2) is displayed on the y-axis. Each dot corresponds to an MII oocyte, and the green and red colours indicate the *Padi6*^+/+^ and *Padi6*^P620A/P620A^ genotype, respectively. (**b**) Volcano plot displaying the differentially expressed genes (DEGs) between *Padi6*^P620A/P620A^ and *Padi6*^+/+^ MII oocytes. The Log2 Fold Change (Log2FC) is reported on the x-axis, and the significance expressed as –log10(FDR). The dashed lines represent the thresholds used for DEGs (|Log2FC| > 1 and FDR < 0.01) on the y-axis. The significant DEGs are indicated in red. (**c**) Heatmap displaying the DEGs between the *Padi6*^P620A/P620A^ and *Padi6*^+/+^ MII oocytes. Columns represent different samples and rows different genes. By Hierarchical clustering we identified three clusters: Cluster 1 (orange), Cluster 2 (red) and Cluster 3 (green). The expression profile is rapresented as scaled FPKM. (**d**) Upset plot intersecting the germinal vesicle marker genes (GVg) with the MII DEGs. The MII.PC genes are obtained from PC1, which separates the C2 cluster in red (**c**). The MII.other genes correspond to the remaining DEGs. (**e**) Barplot showing the top enriched GO terms in the MII DEGs. The y-axis represents the percentage of genes covered in each GO term. (**f-j**) Violin and boxplots displaying the DNA methylation of whole genome divided in 100-CpG tiles (**f**), gene bodies (**g**), promoter regions (TSS-3000+100) (**h**), CpG islands (**i**) and LINEs (**j**) of the *Padi6*^P620A/P620A^ and *Padi6*^+/+^ MII oocytes. In each case, the violin plots represent the merged data from 4 *Padi6*^P620A/P620A^ and 6 *Padi6*^+/+^ MII oocytes.

Whole-genome methylation profiles of the MII oocytes were determined by employing single-cell bisulfite sequencing (scBS-seq). We found some changes in the whole-genome distribution of DNA methylation between the *Padi6*^P620A/P620A^ and *Padi6*^+/+^ oocytes. In particular, a slight hypomethylation was found in gene bodies, promoters, CpG islands (CGIs) and LINEs of the *Padi6*^P620A/P620A^ oocytes (Fig. 5f-j, Supp. Table S4-8), while no significant difference was identified in low complexity regions, LTR elements and simple repeats (Fig. S2a). No significant difference in the methylation of the imprinted germline DMRs was found as well (Fig. S2b,c). In summary, *Padi6*^P620A/P620A^ MII oocytes demonstrated altered RNA profiles with respect to controls, which in part corresponded to elevated abundance of transcripts typical of immature oocyte stages and deregulation of transcripts involved in cytoplasmic functions. Whole-genome DNA methylation was slightly affected but maternal methylation imprinting was unchanged.

### *Padi6*^MatP620A/+^ embryos show failure of ZGA and epigenetic reprogramming

To better investigate the molecular basis of the developmental arrest of the embryos derived from *Padi6*^P620A/P620A^ oocytes, we determined the transcriptome of the 2-cell embryos generated by IVF. This analysis revealed 624 2c-DEGs (FDR < 0.05) between the *Padi6*^MatP620A/+^ and *Padi6*^+/+^ embryos. Among these, 395 were down-regulated and 229 up-regulated (Fig. 6a, Supp. Table S9). To investigate the possible role during maternal to zygotic transition, we first identified the genes that were differentially expressed between the *Padi6*^+/+^ MII oocytes and 2-cell embryos. This analysis revealed 2146 upregulated and 2643 downregulated genes between MII oocytes and 2c embryos that we defined ZGA and Mat Decay genes, respectively (Supp. Table S10). Using these genes as reference, we found that 265 ZGA and 113 Mat Decay genes were deregulated in the *Padi6*^MatP620A/+^ 2-cell embryos. Interestingly, 95% of these ZGA 2c-DEGs were down-regulated and 93% of the Mat Decay 2c-DEGs were up-regulated (Fig. 6b, Supp. Table S11). Notably, 25% of these deregulated ZGA genes were related with the major ZGA, 4% with the minor ZGA and 22% were classified as minor and major ZGA genes (Fig. 6b) ((Wang et al. 2022). In particular, we found normal expression levels of *Dux* family genes, known to be involved in the minor ZGA, and down-regulation of genes strongly associated with the major ZGA, such as *Zscan* family, *Tcstv* family and *Zfp352* (Fig. 6c). To investigate the impact of the 2c-DEGs on embryo development, we performed a STRING k-mean cluster analysis and found that 71 out of the 378 deregulated gene products interacted and could be sorted in 4 clusters associated with different functions: I. histone binding, II. RNA binding, III. regulation of RNA polymerase activity, IV. heterogeneous pathways including leukemia inhibitory factor receptor and inositol triphosphate kinase pathways, indicating a crosstalk between the affected ZGA and Mat decay genes, which may underlie the 2-cell development arrest of the *Padi6*^MatP620A/+^ embryos (Fig. 6d and S3a). As in the MII oocytes, we tested whole genome methylation by scBS-seq. The results obtained demonstrated a global hypermethylation of *Padi6*^MatP620A/+^ compared to *Padi6*^+/+^ embryos (Fig. 6e, Supp. Table S12). This increase involved promoter regions, LINEs, LTRs and Simple repeats (Fig. 6f, S4a, Supp. Table S13-14). In particular, sorting the LINEs on the basis of their evolutionary age (Sookdeo et al. 2013), we found that the younger elements were more strongly hypermethylated than older ones (Fig. 6g and S4b). Moreover, we found a highly significant hypermethylation in the evolutionary younger and more active RLTR4*/*murine leukemia virus (MLV) LTR repeats (Kozak 2014), with respect to other LTR subclasses classified by Repbase Update (Fig. 6h, Supp. Table S15; Jurka 2005). In contrast, only slight and non-significant differences were found in the methylation of the imprinted DMRs (Fig. S4 c,d). Furthermore, comparison of whole-genome gene expression and DNA methylation revealed no significant correlation between differential gene expression and promoter methylation in the *Padi6*^MatP620A/+^ embryos (Fig. S3b). In summary, the *Padi6*^MatP620A/+^ embryos failed to fully activate major ZGA genes and downregulate Mat decay genes at the 2-cell stage and this was accompanied by a genome-wide hypermethylation that severely affected active LINEs and LTRs.

**Figure 6.**
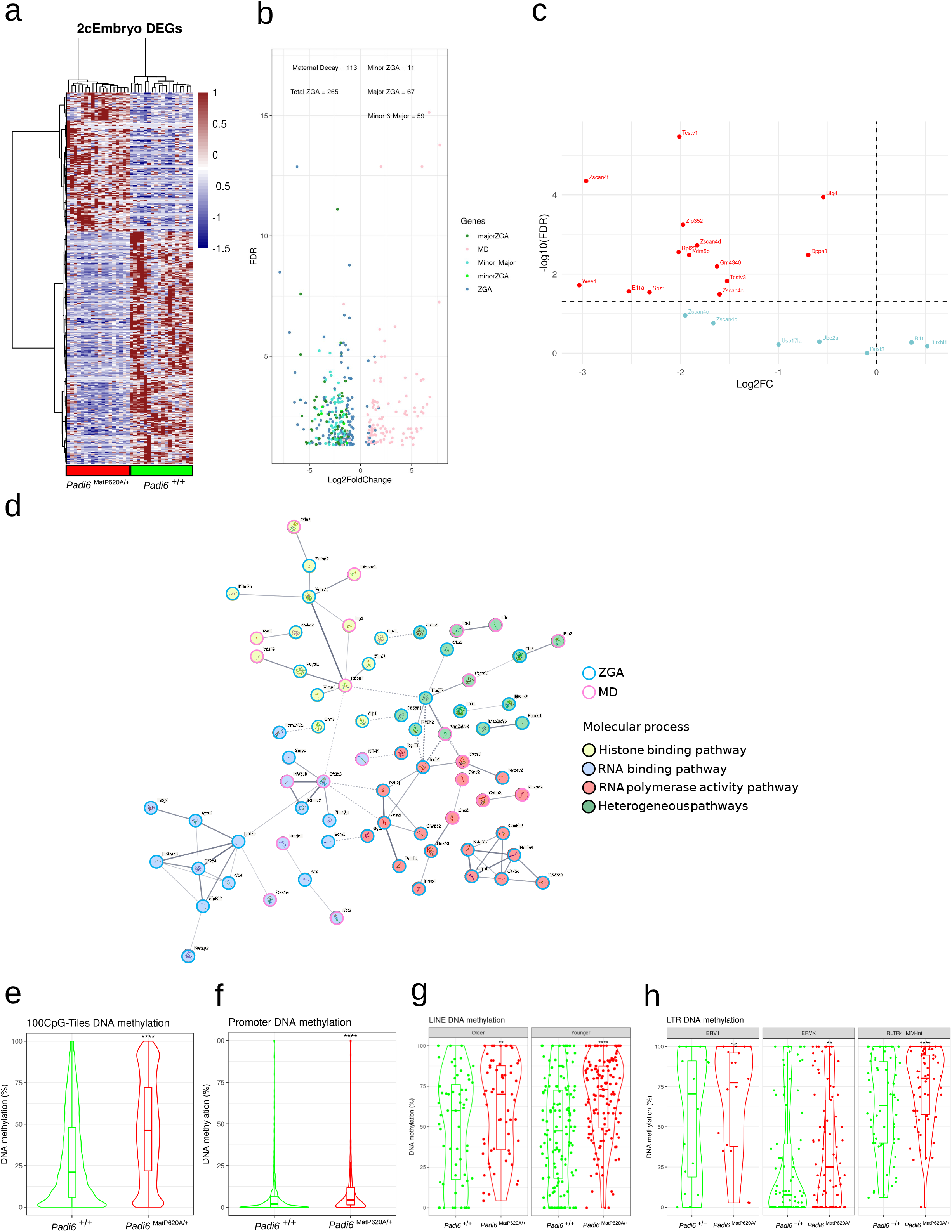
scRNA-seq and BS-seq analyses of *Padi6*^MatP620A/+^ 2-cell embryos. (**a**) Heatmap displaying the 624 DEGs between the *Padi6*^P620A/+^ and *Padi6*^+/+^ 2-cell embryos. Note that only two distinct clusters characterized by the two different genotypes segregate by hierarchical clustering. The expression profile is rapresented as scaled FPKM. (**b**) Volcano plot displaying color-coded DEGs based on their role in ZGA or maternal decay. (**c**) Dotplot showing the differential expression of genes with important roles in ZGA. The Log2 Fold Change (Log2FC) is reported on the x-axis and the significance expressed as –log10(FDR) on the on the y-axis. (**d**) STRING analysis of the proteins encoded by the ZGA and mat decay DEGs. The lines connecting the circles indicate protein-protein interactions. The line thickness reflects the strength of the interaction. The circles corresponding to the ZGA DEGs are depicted with blue borders, the mat decay DEGs with pink borders. The circles are filled with different colours according to the molecular process in which the gene is involved. (**e-h**) Violin and boxplots displaying the DNA methylation of whole genome divided in 100-CpG tiles (**e**), promoter regions (TSS-3000+100) (**f**), Old and Younger LINEs (**g**) and LTR subclasses (**h**) of the *Padi6*^MatP620A/+^ and *Padi6*^+/+^ 2-cell embryos. In each case, the violin plots represent the merged data from 15 *Padi6*^MatP620A/+^ and 14 *Padi6*^+/+^ 2-cell embryo cells.

### The *PADI6***^P620A^** variant is associated with abnormal DNMT1 cellular localization

To investigate the possible cause of the hypermethylation in the *Padi6*^MatP620A/+^ 2-cell embryos, we evaluated the expression levels of the DNA methylation-associated genes *Dnmt1*, *Dnmt3a*, *Tet3* and *Uhrf1* by using the scRNA-seq dataset. No difference was found in expression of these genes between mutant and control MII oocytes as well as between mutant and control 2-cell embryos (Fig. 7a,b). Because DNMT1 activity is controlled through cytoplasm to nucleus transport during early development (Chen and Zhang 2020), we analysed the subcellular localization of the proteins described above by immunofluorescence. We found that DNMT1 had the expected cytoplasmic localization in the *Padi6*^+/+^ GV oocytes, but was mostly concentrated in the nucleus of *Padi6*^P620A/P620A^ GV oocytes. Notably, UHRF1, the DNMT1 accessory protein (Chen and Zhang 2020), also appears to lose its normal cytoplasmic localization in *Padi6*^P620A/P620A^ GV oocytes, but does not appear to become as concentrated in the nucleus (Fig. 7f). In contrast to DNMT1 and UHRF1, the cellular localizations of both DNMT3A and TET3 were unaffected (Fig. 7c-e). A similar result was obtained in the zygotes. *Padi6*^MatP620A/+^ zygotes showed a strong DNMT1 delocalization in both maternal and paternal pronuclei compared to *Padi6*^+/+^ zygotes (Fig. 7g). In contrast, the subcellular localization of DNMT3A and TET3 was not altered (Fig. 7h,i). Taken together, these results demonstrate that DNMT1 was incorrectly localized in the nucleus in both *Padi6*^P620A/P620A^ oocytes and *Padi6*^MatP620A/+^ 2c embryos, providing a possible explanation for the embryo hypermethylation.

**Figure 7.**
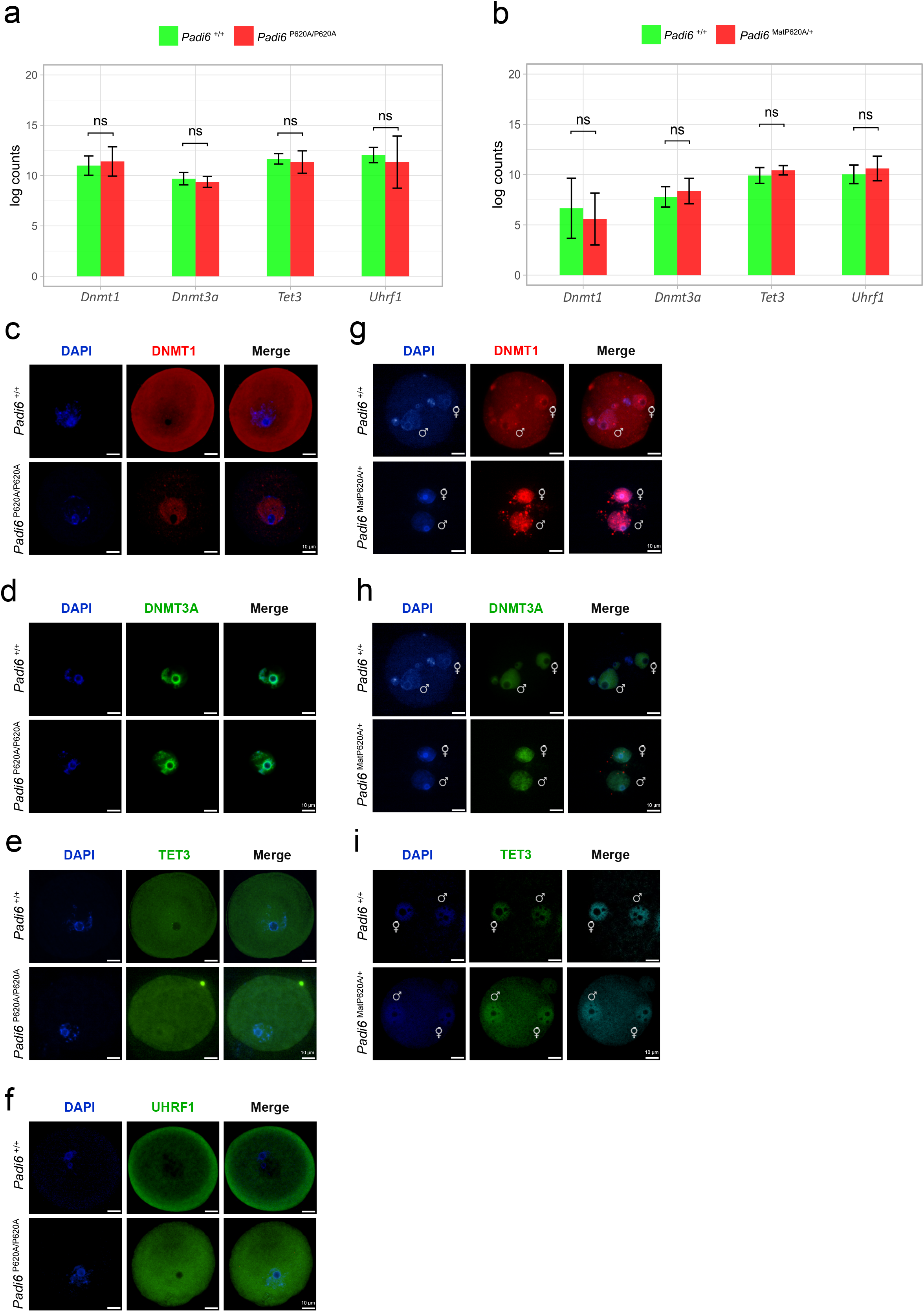
Effect of the *Padi6*^P620A^ variant on expression and subcellular localization of DNMTs, UHRF1 and TET3. (**a**) Barplot representing the expression level of the *Dnmt1*, *Dnmt3a*, *Tet3* and *Uhrf1* genes calculated as log2 mean count of scRNAseq data obtained from 18 *Padi6*^+/+^ (green) and 20 *Padi6*^P620A/P620A^ (red) MII oocytes. (**b**) Barplot representing the expression level of the genes shown in (a) in 18 *Padi6*^+/+^ (green) and 18 *Padi6*^MatP620A/+^ (red) 2-cell embryos. (**c-f**) Representative images of *Padi6*^+/+^ (top) and *Padi6*^P620A/P620A^ (bottom) GV oocytes immunostained with anti-DNMT1 (red) (**c**), –DNMT3A (green) (**d**), –TET3 (green) (**e**) and –UHRF1 (green) (**f**) antibodies. (**g-i**) Representative images of *Padi6*^+/+^ (top) and *Padi6*^MatP620A/+^ (bottom) zygotes immunostained with anti-DNMT1 (red) (**g**), –DNMT3A (green) (**h**), –TET3 (green) (**i**) antibodies. The maternal and paternal pronuclei are indicated by ♀ and ♂, respectively. The data in (**a)** and **(b)** are mean ± s.d. and were analysed using an unpaired two-tailed Student’s *t*-test (ns, *P* < 0.05; \*\**P* < 0.01; \*\*\**P* < 0.001). **c**, **d**, **e**, **f**, **g**, **h**, and **i** are representative images from three independent biological replicates.

## Discussion

The integrity of the oocyte-derived SCMC protein PADI6 is needed for pre-implantation development in humans and mice, and *PADI6* variants are associated with imprinting disorders. In particular, maternal PADI6 is needed for *de novo* transcription and translation in 2-cell embryos (Yurttas et al. 2008). However, the mechanisms by which PADI6 affects these processes in early development are poorly defined. By applying single-cell analysis to a knock-in mouse line in which a BWS-associated *PADI6* variant was modelled, we provided functional insights into the mechanism by which the maternal *Padi6* gene controls epigenetic reprogramming and zygotic genome activation in pre-implantation embryos.

Transcriptome analysis demonstrated down-regulation of many ZGA genes in the *Padi6*^MatP620A/+^ 2-cell embryos. Although both minor ZGA and major ZGA were found to be repressed, we noticed that the early 2c markers *Duxf3* and *Duxbl1* (De Iaco et al. 2017) were normally expressed while the DUX-target genes *Zscan4c*, –*4d* and *-4f, Zfp352* and *Trim43* (Geng et al. 2012); (Sugie et al. 2020) were down-regulated in the *Padi6*^MatP620A/+^ embryos, suggesting that the PADI6 variant mostly affected the major ZGA in the late 2-cell stage. Accordingly, further 2c– and ZGA-specific genes (Genet and Torres-Padilla 2020), such as *Tcstv1* and *Tcstv3*, *Zfp352*, *Elf1a*, *Wee1*, were found down-regulated. Several RNA polymerase genes were down-regulated as well, and this likely prevented the progression of embryogenesis further than the 2-cell stage.

Failure of ZGA was accompanied by a dramatic hypermethylation of the whole genome in *Padi6*^MatP620A/+^ 2-cell embryos. Gain of methylation involved gene promoters, but no correlation was found between gene methylation and expression. Repetitive elements, in particular LTRs and LINEs, were hypermethylated as well. It has been demonstrated that DUX activates transcription of MERVL retrotransposons and that this is required for development beyond the 2-4 cell stage (Hendrickson et al. 2017; Sakashita et al. 2023). In addition, several 2-cell genes, such as *Zscan4*-, *Tcstv1/3*, *Eif1a*, *Tho4* (*Gm4340*) *Tdpoz*-, and *Zfp352*, which we found downregulated in the *Padi6*^MatP620A/+^ embryos, are transcribed as chimeric genes with MERVL LTRs (Macfarlan et al. 2012). Concerning the LINEs, it has been shown that their activation at the 2-cell stage regulates global chromatin accessibility and their targeted repression strongly affects developmental rate (Jachowicz et al. 2017). Interestingly, we found that the more recently evolved LTR and LINE elements that retain transcription had the most highly significant gain of methylation. DNA methylation is required to maintain repression of ERV and LINE transposons in post-implantation embryos (Dahlet et al. 2020) (Graham-Paquin et al. 2023). These results suggest that hypermethylation of the LTR and LINE elements likely interfering with their timely activation in early embryos has a major role in the ZGA failure of *Padi6*^MatP620A/+^ embryos.

In preimplantation embryos, DNMT1 and its cofactor UHRF1 are actively retained in the cytoplasm, thus enabling demethylation of the genome by passive dilution at DNA replication (Cardoso and Leonhardt 1999) (Maenohara et al. 2017). In contrast, we found that DNMT1 is mostly concentrated in the pronuclei of the *Padi6*^MatP620A/+^ zygotes. The aberrant nuclear localization of DNMT1 and UHRF1 is already evident in *Padi6*^P620A/P620A^ GV oocytes, suggesting that this is a cause rather than the consequence of the impaired ZGA of the mutant 2-cell embryos. We expect that the percentage of symmetrically methylated CpGs increases during replication of the zygotic DNA in the presence of nuclear DNMT1 activity and this could contribute to ZGA failure in the *Padi6*^MatP620A/+^ embryos. We did not find DNA hypermethylation in the *Padi6*^P620A/P620A^ MII oocytes, despite the strong nuclear staining of DNMT1, but perhaps the apparently more limited nuclear presence of UHRF1 prevents excessive methylation. The mechanism by which PADI6 controls intracellular DNMT1 localization remains elusive. However, DNMT1 was also found in the nuclei of *Nlrp2*-, *Nlrp14*– and *Pgc7/Stella* maternal-knockout embryos, which all displayed cytoplasmic defects (Mahadevan et al. 2017; Li et al. 2018; Meng et al. 2023; Yan et al. 2023). Furthermore, *Nlrp14*– and *Pgc7/Stella* Mat-knock-out embryos also displayed genomic hypermethylation and ZGA failure. Interestingly, DNMT1 shows a SCMC-like localization in mouse oocytes (Mahadevan et al. 2017) and Fig. 7c), suggesting that the SCMC may contribute to its retention in the cytoplasm at this developmental stage.

After fertilization, the zygotic genome is inert and mRNAs derived from the oocyte are translated to direct embryo development up to the 2-cell stage (Chen et al. 2023). Oocyte transcripts are progressively decayed through either maternal (M-decay) and zygotic (Z-decay) pathways (Jiang and Fan 2022; Chen et al. 2023). In particular, the Z-decay pathway involves the polyA-binding proteins PABPC1 and PABPN1, both of which were found down-regulated at the transcriptional level in *Padi6*^MatP620A/+^ embryos, suggesting that the up-regulation of Mat-decay genes is a consequence of the ZGA failure. Additional factors, such as *Btg4* (Zheng et al. 2020) that is slightly down-regulated, may be involved in the deregulation of Z-decay in the *Padi6*^MatP620A/+^ embryos.

Although no significant difference in ovulation efficiency was observed between the *Padi6*^P620A/P620A^ and the *Padi6*^+/+^ mouse lines, the mutant MII oocytes displayed evidence of maturation defects with an increase of cells characterized by abundance of GV-specific markers and genes involved in cytoplasmic and metabolic processes. For instance, we found upregulated genes like *Idh2*, *Mogs*, *Gpd1*, *Decr1*, *Alox5ap*, and *Atp2b2,* which stimulate energy production, glycosylation, lipid metabolism, and calcium signalling. These molecular changes are crucial for the intricate process of oocyte development, ensuring successful fertilization and embryonic development. PADI6 also interacts with the maternal-effect Zygotic arrest-1 (ZAR1) protein that is involved in oocyte maturation by interacting with cytoplasmic lattices and controlling mRNA stability and translation (Rong et al. 2019). Thus, our data are consistent with a role of PADI6 in the control of oocyte maturation. It should also be taken into account that in our study the MII oocytes were collected after PMSG+hCG stimulation that may have pushed immature oocytes to ovulate (Moor et al. 1985) and enhanced the difference between the mutant and the control oocytes.

We observed that the *Padi6*^P620A/P620A^ oocytes correctly transcribed the *Padi6* gene, but failed to accumulate the PADI6 protein above the level that was detectable by western blotting and immunofluorescence. This was unexpected because the human variant *PADI6*^P632A^ was predicted to be deleterious by SIFT and possibly damaging by Polyphen, but stabilizing by the SDM program (Cubellis et al. 2020). We reanalysed the impact of the orthologous human and mouse variants on protein stability by using the more recent and integrated computational method Dynamut (Rodrigues et al. 2018) and two further structure-based prediction tools (mCSM and DUET) (Pires et al. 2014b; Pires et al. 2014a) and found that in this case both variants were predicted to be destabilizing (Suppl. Table S16). Thus, we consider this prediction more reliable on the basis of the results obtained *in vivo*. Nevertheless, a woman carrying the *PADI6*^P632A^ variant in compound heterozygosity with a null mutation conceived two children with BWS and MLID and did not report any fertility problem (Cubellis et al. 2020). So, the phenotype associated with the mouse *Padi6*^P620A^ variant is more severe than that associated with the orthologous human variant. In addition, we did not find any significant change in the methylation of the imprinted DMRs in either *Padi6*^P620A/P620A^ MII oocytes or *Padi6*^MatP620A/+^ 2-cell embryos. This is consistent with the findings obtained with the *Nlrp2* and *Nlrp14* knock-out mouse lines, in which minor and no imprinting defects were reported (Mahadevan et al. 2017; Yan et al. 2023). In addition, the normal methylation profiles of the *Padi6*^P620A/P620A^ MII oocytes indicates that at least in the mouse PADI6 is not required for *de novo* methylation as demonstrated for human KHDC3L (Demond et al. 2019). Some redundancy in the human SCMC proteins (e.g., NLRP7 is absent in mice) or differences in the ZGA length may explain these phenotypic differences between humans and mice. Also, it is possible that the embryos surviving the impaired ZGA may develop defects in maintenance of imprinted methylation after the 2-cell stage.

In conclusion, our study demonstrates that the maternal-effect gene *Padi6* has a major role in controlling epigenetic reprogramming and zygotic genome activation in early mouse embryos and is consistent with a model in which the SCMC is necessary for cytoplasmic localization of DNMT1 that in turn is crucial for the genome-wide DNA demethylation occurring during the maternal to zygotic transition.

## Materials and Methods

### Generation of *Padi6*^P620A^ mouse line

The P620A variant was introduced into the *Padi6* locus of E14Tg2a4 (129/SV) ESCs by homologous recombination. Homology arms obtained from a BAC clone containing Padi6 isogenic DNA were inserted into a pFLRT3D vector modified for Flp/Frt-based recombination. Details of the targeting strategy are available upon request. The linearised targeting vector was electroporated into ESCs and, upon G418 selection, three positive clones were identified by PCR and confirmed by Southern blotting with one internal and two external probes. The presence of the mutation was assessed by allele specific PCR. In order to generate chimeras, ten to fifteen ESCs from one clone were microinjected into C57Bl/6 blastocysts and ten blastocysts were reimplanted in each uterine horn of pseudopregnant B6D2 females, anaesthetised with 80 mg/kg Ketamine and 10 mg/kg Xylazine. Highly chimeric animals were crossed to Flp recombinase deleter mice to remove the neomycin cassette. The offspring were used to establish the *Padi6*^P620A^ mouse colony. All experimental procedures involving animals were conducted according to the authorization n° 830/2020-PR, released by the Italian Ministry of Health. The primers used are listed in Supp. Table S17.

### Mouse genotyping

The presence of the *Padi6* variant, the removal of Neo box and the presence of the flippase gene were demonstrated by PCR on tail DNA extracted with standard procedures. Amplicons of 430 bp and 290 bp were amplified from the mutant or wild type *Padi6* alleles, respectively, by using the primers Padi6_RimF and Padi6_RimR. This difference was due to a fragment of the NEO box remaining after the flippase recombination. The presence of the mutation was also identified by digestion with the restriction enzyme StuI, which was generated by the mutation, after amplification of a 420 bp fragment with the primers For LA HindIII + SnaBI and Rev GEN Padi6. The presence of the NEO box was detected using the primers PGK1A For and Padi6_RimR. The presence of the flippase was detected using the primers FLP_F and FLP_R. The primer sequences are listed in the Supp. Table S17.

### Assessment of fertility

2-4 month-old female *Padi6*^+/+^, *Padi6*^+/P620A^ or *Padi6*^P620A/P620A^ mice were housed with >8 week-old *Padi6*^+/+^ male mice at 4-6pm and checked for the presence of vaginal plug the next morning. Positive females were monitored for pregnancy and delivery.

### MII oocyte collection and *in vitro* fertilisation

Heterologous in vitro fertilisation was performed according to the method described by Taft (2017). 8-12 week-old males and 4-8 week-old females were used. To induce superovulation, the females received intraperitoneal injections of pregnant mare serum gonadotropin (PMSG) on day 1 at 2 pm and human chorionic gonadotropin (hCG) after 52 hours. Males were sacrificed approximately 12 hours after the female hCG injection, cauda epididymis and vasa deferentia were dissected and spermatozoa were collected in human tubal fluid medium (HTF) (EmbryoMax® HTF MR-070-D). Spermatozoa were counted and incubated at 37°C within a humidified atmosphere with 5% CO2. Females were sacrificed 13 hours after the hCG injection. Ampullae were placed in 1mg/ml hyaluronidase (H6254-500MG) drops to break down the oocyte-cumulus complex and collect MII oocyte. The oocytes were moved to the HTF medium and 2×10^6^ sperm/ml spermatozoa were added. After 4 – 6 hours of incubation at 37°C, the fertilised oocytes were washed, to remove the spermatozoa e debris, transferred in KSOM medium and incubated overnight at 37°C. The embryos’ status was evaluated the next morning and the following 120 hours.

### DNA and RNA isolation for single-cell genomic analysis

MII oocytes and 2-cell embryos were individually lysed and flash-frozen in 5 μl RLT Plus buffer (Qiagen) and stored at −80°C until further use. For 2-cell embryos, the zona pellucida was removed with Tyrode’s solution (SIGMA-ALDRICH), blastomeres were separated and polar body discarded. DNA and RNA were separated using the G&T protocol (Angermueller et al. 2016). Magnetic beads (MyOne C1, Life Technologies) were washed and annealed to oligo dTs and then used to capture polyadenylated mRNA from individual cell lysates. The supernatant containing the DNA was transferred to a new tube and the beads washed three times in 1xFSS buffer (Superscript II, Invitrogen), 10 mM DTT, 0.005% Tween-20 (Sigma) and 0.4 U/μl of RNAsin (Promega) to remove all DNA residues. Each washing solution was added to the DNA tube to maximise recovery.

### Construction of scBS-seq libraries

The scBS-seq libraries were generated as described (Clark et al., 2017). The DNA was purified from the lysated using a 0.9:1 ratio of Ampure XP Beads (Beckman Coulter) and eluted into 10 μl of water. DNA purified from single cells was treated with the EZ Methylation Direct Kit (Zymo) for bisulfite-conversion following the manufacturer’s instructions. First-strand synthesis was performed in five rounds. Initially, the bisulfite-treated DNA was mixed with 40 μl first-strand synthesis mastermix (1 × Blue Buffer (Enzymatics), 0.4 mM dNTP mix (Roche), 0.4 μM 6NF oligo (IDT)) and heated to 65 °C for 2 min and cooled on ice, followed by the addition of 50U of Klenow exo– and incubation at 37 °C for 30 minutes after slowly ramping from 4 °C. This process was repeated four more times with additional reaction mixture added each time, and the final round was incubated for 90 minutes at 37 °C. Exonuclease digestion was carried out by adding 20U of exonuclease I (NEB) in a total volume of 100 μl, at 37 °C for 1 h. The resulting samples were purified using AMPure XP beads with a 0.8:1 ratio. The beads were mixed with 50 μl second-strand master mix (1× Blue Buffer (Enzymatics), 0.4 mM dNTP mix (Roche), 0.4 μM 6NF oligo (IDT). The mixture was heated to 98 °C for 1 min and cooled on ice. Then, 50U of Klenow exo-(Enzymatics) were added. The mixture was incubated on a thermocycler at 37 °C for 90 min after slowly ramping from 4 °C. The resulting samples were purified using a 0.8:1 ratio of AMPure XP beads, and libraries were amplified using a 50 μl PCR mastermix (1× KAPA HiFi Readymix, 0.2 μM PE1.0 primer, 0.2 μM iTAG index primer) The amplification process involved a 2 min step at 95 °C, followed by 14 cycles of 80 s at 94 °C, 30 s at 65 °C, and 30 s at 72 °C, with a final extension for 3 min at 72 °C. Finally, the scBS-seq libraries were subjected to purification using a 0.7:1 ratio of AMPure XP beads. After purification, the libraries were eluted in 15 µl of water, pooled together, and sequenced. The sequencing process involved the generation of pools of 48 libraries, which were sequenced on an Illumina HiSeq 2500 platform. On average, the libraries were sequenced to generate 13.0 million paired-end reads with a read-length of 75 bp.

### Construction of scRNA-seq libraries

The mRNA bound to the beads was processed further for cDNA conversion. This involved the resuspension of the beads in 10 μl of reverse transcriptase mastermix (100 U SuperScript II (Invitrogen), 10 U RNAsin (Promega), 1 × Superscript II First-Strand Buffer, 5 mM DTT (Invitrogen), 1 M betaine (Sigma), 9 mM MgCl2 (Invitrogen), 1 μM Template-Switching Oligo (TSO, Eurogentec), 1 mM dNTP mix (Roche). The mRNA mixture was reverse transcribed by incubation for 60 min at 42 °C followed by 30 min at 50 °C and 10 min at 60 °C. The cDNA obtained was subjected to PCR amplification by adding 11 μl of 2x KAPA HiFi HotStart ReadyMix and 1 μl of ISPCR primer (2 μM). The amplification was carried out in a thermocycler at 98 °C for 3 min, followed by 15 cycles of 98 °C for 15 s, 67 °C for 20 s, 72 °C for 6 min and the final extension step at 72 °C for 5 min. The amplified product was purified using Ampure XP beads with a 1:1 ratio and eluted into 20 μl of water. Libraries were prepared from 100 to 400 pg of cDNA using the Nextera XT Kit (Illumina) and following the manufacturer’s instructions but with one-fifth volumes. All 96 single-cell RNA-seq libraries were pooled together and sequenced on an Illumina NextSeq platform to an average depth of 4.2 million reads, using paired-end 75 bp read-length settings.

### Histological analysis

Ovaries were collected from 2-4 months old mice and fixed in formalin overnight, dehydrated through graded alcohol solutions and immersed in xylene for 1 h for the diaphanization. They were finally embedded in paraffin blocks and cut with a rotary microtome (Leica RM2235, Germany) into 7-µm thick sections, which were mounted on glass slides. They were de-paraffinized in xylene, rehydrated through graded alcohol solutions and stained with hematoxylin and eosin (H&E). The H&E images of the ovaries were captured by using a Nikon Motorized Optical Microscope.

### Immunostaining of mouse tissues, oocytes and zygotes

GV oocytes and zygotes were fixed with 4% paraformaldehyde in PBS at room temperature for 30 min and washed with PBS three times. Fixed cells were incubated in the permeabilization solution (0.5 % Triton-X100 and 0.05% Tween-20 in PBS) for 30 min. After incubation with a blocking solution (0.1 % BSA and 0.05% Tween-20 in PBS) at room temperature for 1 h, the cells were incubated at 4°C overnight with the primary antibody. After 3x wash, the cells were incubated with the secondary antibody at room temperature for 1 h. Ovarian tissue sections were processed as described for the histological analysis, until de-paraffinization and rehydration. Antigen retrieval was performed by microwave processing at 700 W in 0.01 M (pH 6.0) citrate buffer for 10 min. Slides were incubated with Blocking Solution (0.5% Milk, 10% FBS, 1% BSA in H_2_O, 0.02% NaN3) for 60 min at room temperature in a humidified chamber and the same blocking solution was used to dilute primary and secondary antibodies. The primary antibody was incubated in humidified conditions overnight at 4°C. Secondary antibody was incubated for 1 h at room temperature. Cells and tissues were mounted in SlowFade^TM^ Gold antifade reagent with DAPI (S36938) and observed using Nikon Motorized Optical Microscope with a 20× ocular lens. The primary antibodies used in this study were as follows: anti-PADI6 1:500 (generously provided by Dr. Scott Coonrod, Cornell University, Ithaca, New York); anti-OOEP 1:200 (PA586033); anti-DNMT1 1:250 (Ab13537); anti-DNMT3A 1:500 (Ab188470); anti-TET3 1:400 (NBP2-20602). The secondary antibodies used in this study were as follows: anti-Mouse Alexa Fluor™ 594 1:1000 (A11032); anti-Rabbit Alexa Fluor™ 448 1:500 (A11008); anti-Guinea pig Alexa Fluor™ 594 1:500 (A11076).

### RNA analysis

RNA was isolated from ovaries using the TRI reagent (Sigma-Aldrich) and following the manufacturer’s instructions. Approximately 900 ng of total RNA was retro-transcribed by using the QuantiTect Reverse Transcription Kit (Qiagen) according to the manufacturer protocol. For locus-specific gene expression analysis, cDNA was amplified by real-time PCR using SYBR Green PCR Master Mix (Bio-Rad) on a CFX Connect Real-Time PCR Detection System. The primers used are listed in the Supp. Table S17.

### Western blotting

Tissues were resuspended in RIPA buffer (50 mM Tris HCl pH 7.5, 150 mM NaCl, 1mM EDTA, 1% Triton X100), homogenised with the tissue lyser and incubated for 30 min at 4°C. Proteins were quantified by Quick Start™ Bradford Protein Assay (Bio-Rad). The proteins for western blotting were boiled at 95°C for 10 min in Laemmli buffer and resolved by 8% acrylamide gel. Samples were transferred onto Nitrocellulose membranes (Biorad Transblot) and processed by using standard procedures. Signals were visualised using an ECL method. The Imagej software was used to quantify the bands. The primary antibodies used were: anti-PADI6 1:1000; anti-NLRP5 1:500 (sc514988); anti-TLE6 1:1000 (sc515065). The secondary antibodies were: anti-Mouse 1:10000 (Ab6728); anti-Rabbit 1:10000 (Ab6721); anti-Guinea pig 1:5000 (Ab6908).

### Processing of scRNA-seq data

A total of 96 libraries were adapted and quality trimmed (Phred score <20) using Trim Galore version 0.4.4 (https://www.bioinformatics.babraham.ac.uk/projects/trim_galore/). We also sequenced 4 negative controls to archive technical problems. The good quality reads were aligned to the Genome Reference Consortium mouse genome build 39 (GRCm39) with HiSat2 version 2.1.0 (Kim et al. 2015) in paired-end mode. The data were quantified in SeqMonk and analysed in Rstudio (Supp. Table S18-19). The count matrix was used to calculate the differential expression following the DESeq2 pipeline (Love et al. 2014). The list of ZGA and maternal decay genes was imported into the STRING database (Jensen et al. 2009). Cluster analysis was performed using the “kmean clustering” option with “number of clusters” parameter set to 4. The edges between clusters were set as “Dotted line”. To explore the processes in which those proteins are involved, we (Kolberg et al. 2020) with default options.

### Processing of scBS-seq data

A total of 96 scBS-seq libraries were processed for analysis. The first 6 bp containing the N portion of the random primers, adapters and bases called with poor quality (Phred score <20) were removed using Trim Galore version 0.4.4 (https://www.bioinformatics.babraham.ac.uk/projects/trim_galore/) in single-end mode. Only the good quality reads were used for the alignment to the Genome Reference Consortium mouse genome build 39 (GRCm39) using Bismark version 0.18.2 (Krueger and Andrews 2011) with single-end and non-directional mode followed by deduplication and methylation calling using Bismark functions. The MII scBS-seq libraries having a mapping efficiency <10% or fewer than 500,000 CpGs covered were removed (Supp. Table S20-21). To investigate a possible somatic cell DNA contamination in MII oocytes, we compared the global CpG methylation rates with the methylation of X-chromosome CpG islands, removing the samples with X-chromosome CpG island mean methylation > 12 % (Fig. S5a-c) (Castillo-Fernandez et al. 2020); there was no difference in the proportion of oocytes excluded on this basis between *Padi6*^P620A/P620A^ and control. In total, out of 44 MII oocytes, we kept 10 cells for downstream analysis. The embryos scBS-seq libraries having fewer than 100,000 reads were removed. In total, out of 44 embryos, we kept for 29 the downstream analysis (Fig. S5d,e). The methylation profile for each feature was calculated by the Bisulfite methylation pipeline implemented in SeqMonk. To assess the global methylation differences among the mutant and the wild-type cells we calculated 100-CpG tiles defined using SeqMonk. The CpGI, the Rmsk and the VISTA enhancers tracks were downloaded from UCSC Genome Browser and converted to mm39 coordinates by liftOver tool. The gene body, the promoter (gene Upstream –3000+100) and the other feature methylation were quantitated by the Bisulfite pipeline in Seqmonk. Coordinates for imprinted DMRs were used to quantitate the methylation in both MII and embryos (Supp. Table S22).

### Whole-genome sequencing

Whole-genome sequencing was performed on genomic DNA extracted from the tail of a homozygous *Padi6*^P620A/P620A^ female. The DNA samples were sequenced 150 bp pair-end at BIODIVERSA srl Service (Milan, Italy) using the Nextera DNA Prep illumina and the Illumina NovaSeq6000 platform. The bioinformatic analysis was performed as previously reported. The read quality was checked using the FASTQC tools (http://www.bioinformatics.babraham.ac.uk/projects/fastqc/). Reads were aligned to the mouse genome reference assembly (mm39) using the BWAmem software package v0.7.17 (Li and Durbin 2009). PCR duplicates were filtered out by Picard v2.9 (http://picard.source forge.net), and the Dellytools v1.1.6 (Rausch et al. 2012) with default parameters were employed to calculate the structural and CNV variants.

### Competing Interest Statement

The authors declare no competing interests.

## Acknowledgments

We thank Scott A. Coonrod for the generous gift of the anti-PADI6 antibody and Megan Hamilton of the Babraham Genomics Facility for support with sequencing.This work was supported by the grant PNRR-MR1-2022-12376622 funded by the Italian Ministry of Health (to AR and CG). Work in G.K.’s lab was funded by the UK Biotechnology and Biological Sciences Research Council (BBS/E/B/000C0423) and Medical Research Council (MR/K011332/1, MR/S000437/1).

## Author Contribution

Author contributions: C.G., Fr.C., D.A., Fl.C., S.C., G.K, and A.R. designed experiments and interpreted results. C.G., L.A., D.A., G.R. and S.C. performed experiments. Fr.Ce., C.G., B.H.M., MV.C, A.G. and S.A. performed bioinformatic analyses. C.G., Fr.C. and A.R. wrote the manuscript with input from all authors.

## Availability of data and materials

Raw data supporting the findings of this study have been deposited under accession code GSE… in the Gene Expression Omnibus repository.

